# Expansion of novel biosynthetic gene clusters from diverse environments using SanntiS

**DOI:** 10.1101/2023.05.23.540769

**Authors:** Santiago Sanchez, Joel D. Rogers, Alexander B. Rogers, Maaly Nassar, Johanna McEntyre, Martin Welch, Florian Hollfelder, Robert D. Finn

## Abstract

Natural products biosynthesised by microbes are an important component of the pharmacopeia with a vast array of biomedical and industrial applications, in addition to their key role in mediating many ecological interactions. One approach for the discovery of these metabolites is the identification of biosynthetic gene clusters (BGCs), genomic units which encode the molecular machinery required for producing the natural product. Genome mining has revolutionised the discovery of BGCs, yet metagenomic assemblies represent a largely untapped source of natural products. The imbalanced distribution of BGC classes in existing databases restricts the generalisation of detection patterns and limits the ability of mining methods to recognise a broader spectrum of BGCs. This problem is further intensified in metagenomic datasets, where BGC genes may be incomplete. This work presents SanntiS, a new machine learning-based tool for identifying BGCs. SanntiS achieved high precision and recall in both genomic and metagenomic datasets, effectively capturing a broad range of BGCs. Application of SanntiS to MGnify metagenomic assemblies led to a resource containing 1.9 million BGC predictions with associated contextual data from diverse biomes and demonstrates a significant fraction of novelty compared to equivalent isolate genomes datasets. Subsequent experimental validation of a novel antimicrobial peptide detected solely by SanntiS, further demonstrates the potential of this approach for uncovering novel bioactive compounds.

## Introduction

The microbial world harbours an enormous reservoir of natural products (also termed secondary metabolites), many of which are widely studied and exploited as therapeutic molecules in a vast range of medical applications (1–3), especially in the area of antimicrobials. Many natural products have industrial applications, including food preservation (4), animal growth promoters (5), and phytostimulants (6). Studies on the role of natural products in their native environments suggest they play a major part in many ecological processes due to their wide range of functions such as microbial defence systems, signalling molecules for biofilm formation, or long-distance mediation between bacteria and their surrounding neighbours (7).

The era of genomics has revolutionised approaches for detecting biosynthetic gene clusters (BGCs), which are sets of colocated genes that encode the molecular machinery necessary for producing natural products. Classically, molecular biology techniques such as gene knockout have been used to identify BGCs, but these methods are both time- and resource-intensive. Today, comparative genomics has emerged as a powerful tool for identification and systematic evaluation of BGCs, greatly increasing the discovery rate by reducing both time and cost. Despite these advancements, the discovery of BGCs remains limited by our ability to cultivate and isolate microorganisms for genome sequencing, a feature often exacerbated when organisms are found living in complex environments.

Metagenomics provides a cultivation-independent approach for understanding the taxonomic and functional compositions of microbial communities, and has provided new insights into their complex dynamics and their relationship to their environment. The past five years have witnessed an increasing availability of deeply sequenced short-read metagenomic datasets, which has enabled the assembly of contigs capable of spanning BGCs. This, coupled with emerging long-read metagenomic sequence datasets, provides a rich resource for identifying BGCs and understanding their heterogeneity and distribution in different environments. The study of natural products and the BGCs that give rise to their synthesis at the microbial-community level has underlined their potential as a source of molecular novelty (8, 9), the importance of chemical interactions in shaping these communities (10), and highlighted their vast potential applications (11).

BGCs are commonly classified based on the chemical structure of the natural product and/or the characteristics of the biosynthetic pathway. Broad classes such as Alkaloids, Non-Ribosomal Peptides (NRPs), Polyketides, Ribosomally synthesised and Post-translationally modified Peptides (RiPPs), Saccharides or Terpenes are widely used for classification (12), with each class being subdivided based on specific characteristics. Some computational approaches to detecting BGCs have focused on detecting specific classes and/or subclasses of BGC (13–15), while others have aimed to predict BGCs from all classes (16–19). Regardless, a key challenge persists: the generalisation of detection patterns across a diverse array of BGCs. This challenge is confounded by the significant imbalance of experimentally validated BGC classes in established databases, such as MIBiG (20) (Supplementary Figure 1b), which limits the ability to identify a wide spectrum of BGCs. Fragmented assemblies, as typically produced from metagenomic assembly, further exacerbate this limitation as BGC genes may be distributed across multiple contigs. Consequently, current methods have difficulties in accurately detecting BGCs in such datasets, highlighting the need for a more robust and adaptable approach. In this work, we address the aforementioned challenges associated with current BGC prediction through the development of a novel machine learning-based approach that is suitable for both genomic and metagenomic datasets, and capable of detecting a broad range of BGCs. We demonstrate the potential of this method through extensive benchmarking and experimental validation of a natural product that was not detected by any of the compared methods. Applying SanntiS to 33, 924 metagenomic assemblies from a wide array of biomes, we created a resource of BGC predictions with contextual data, providing new insights into the diversity and distribution of BGCs found in various environments.

## Results and Discussion

### A machine learning approach for BGC detection

Several widely used methods are currently available for prediction of BGCs in genomic datasets, including ClusterFinder, antiSMASH, deepBGC, and GECCO (16–19). Our benchmarking exercises (see section below) show that the ability of these methods to detect certain classes of BGCs is limited or impacted by high false positive rates. While the machine learning-based methods ClusterFinder and deepBGC have high recall, it comes at the expense of precision. To address such limitations, we developed a BGC detection method based on deep learning: SanntiS (**S**econdary metabolite gene cluster **a**nnotations using **n**eural **n**etworks **t**rained on **I**nterPro **s**ignatures; Supplementary Figure 1a). SanntiS was trained using a positive set of 1, 858 non-redundant experimentally validated BGCs, collected from MiBIG DB v2.0 (20) and a negative set of 5, 574 randomly selected nucleotide regions from the species representative genomes of GTDB (release 86 (23)).

**Figure 1:**
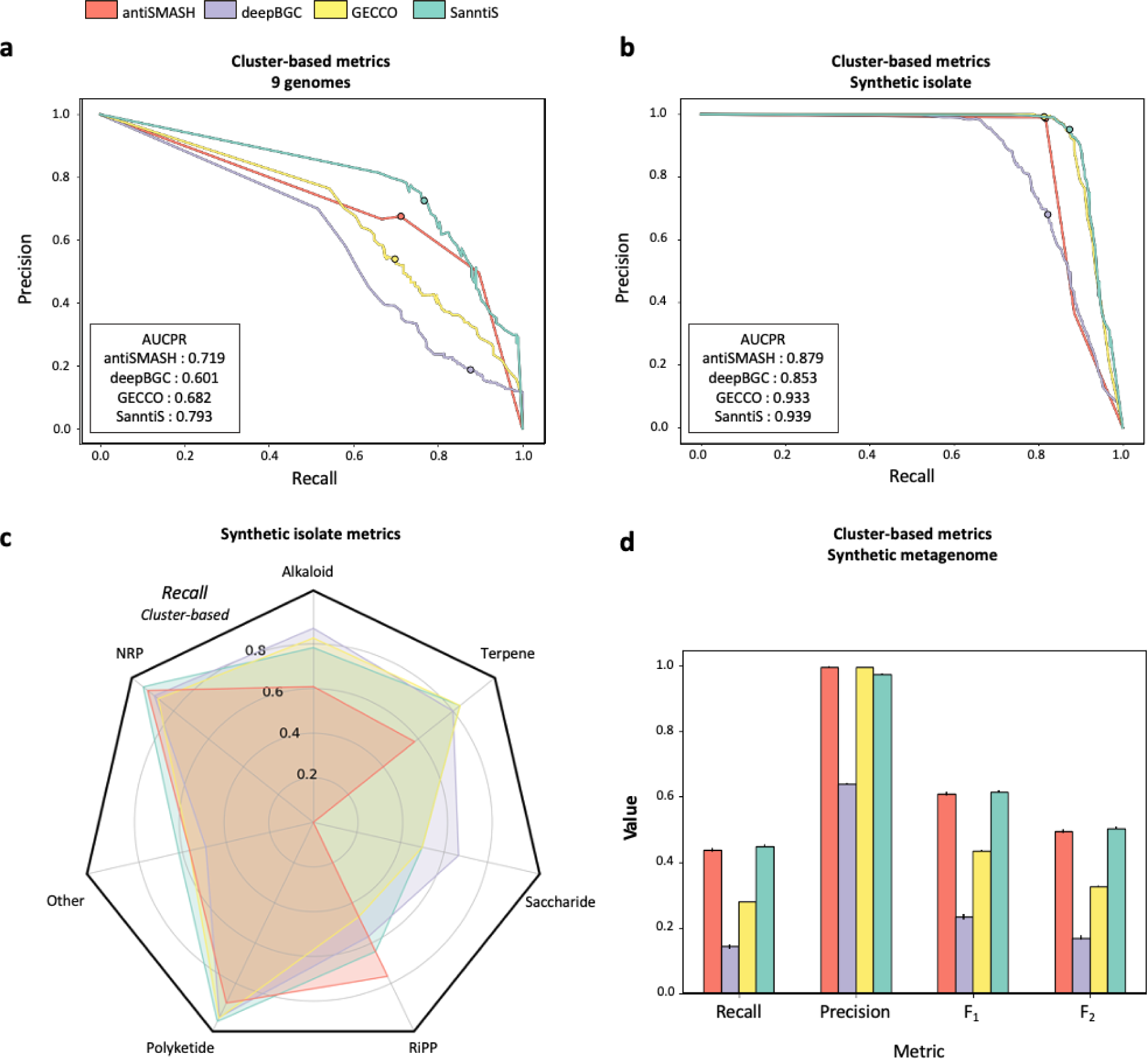
SanntiS performance on genomic and metagenomic datasets with comparison to antiSMASH, deepBGC, and GECCO. (a) Cluster-based precision-recall curve of each method at different probability thresholds in 9 genomes. (b) Cluster-based recall, precision, F1 and F2 scores of each method using default parameters in 9 genomes. (c) Cluster-based recall for each BGC class in a synthetic isolate dataset. The following correspond to the number of BGCs in each class and the percent of eukaryotic BGCs in brackets: Alkaloid, n=23 (31%); NRP, n=233 (23%); Other, n=81 (13%); Polyketide, n=187 (47%); RiPP, n=81 (4%); Saccharide, n=6 (50%); Terpene, n=50 (93%). (d) Cluster-based recall, precision, F1 and F2 scores and standard deviation bars of each method in the 3 replicates of synthetic metagenomic assembly.

At the core of SanntiS is the detection model, an artificial neural network (ANN) with a one-dimensional convolutional layer, plus a bidirectional long short-term memory (BiLSTM). The model was developed using linearized sequences of protein annotations based on a subset of InterPro (see Methods) as input. The use of InterPro resulted in a 13% increase in the number of annotated proteins in BGCs (Supplementary Figure 1b) when compared with using Pfam (34) alone, which is the functional annotation method commonly used by other BGC detection methods. The detection model was further optimised to address the challenge of predicting BGC classes that have a limited representation in the training set. To achieve this, we employed a duration robust loss function (RLF, orange in Supplementary Figure 1c; (35)). RLF mitigates the issue of class imbalance, which can arise from the disparities in BGC counts by class and the variation in the duration of detection events – in this context, the disparities in length across different BGC classes. It accomplishes this by adjusting the binary-cross entropy loss function based on the complexity of detecting a specific event. Specifically, this results in an increase in the overall performance of the model (Supplementary Figure 1c), from an area under the precision-recall curve (AUCPR) of 0.60 to 0.64, when compared with the use of the standard binary cross-entropy (BCE, blue in Supplementary Figure 1c) loss function. Finally, a post-processing random forest (RF) classifier sorts the regions with a high probability of being BGCs (default probability of 0.85) as BGC or non-BGC.

### High precision and recall scores in genomic and metagenomic datasets

We compared the predictive power of our BGC-detection method, SanntiS, with three other methods with similar goals (antiSMASH v6.1.1, deepBGC v0.1.30, and GECCO v0.9.8) using the metrics: precision; recall; F1; and F2 scores. Our assessment of the predictions focused on three key aspects. Firstly, the use of cluster and average metrics. This approach was specifically chosen to mitigate the limitations of point-based metrics in representing range events – a term referring to events that span a sequence of consecutive time points or locations. This is pertinent in situations where the datasets are class-unbalanced and the length distributions of BGCs are uneven. In the cluster approach, the overlap of a predicted region with the ground truth BGC is considered as a true positive (TP) and a predicted region not overlapping any ground truth BGC regions is a false positive (FP; Supplementary Figure 2a). In the average-based approach, precision is calculated for each predicted BGC by dividing the number of TP coding sequences (CDSs) by the total CDSs in the prediction, followed by averaging these values across all predicted regions. Similarly, recall is computed for each ground truth BGC by dividing the number of TP CDSs by the CDSs in the BGC, and then averaging the results across all predicted regions (Supplementary Figure 2b). The second key aspect of our assessment consisted of statistics for individual BGC classes to prevent biases in the results towards the most prevalent classes (see Methods); and, finally, evaluation on both isolate genomes and metagenomics datasets.

To construct our benchmark, we used three distinct datasets: (i) the widely utilised, manually annotated “9 genomes ” for BGC regions; (ii) a synthetic isolate genome, constructed by combining manually annotated BGCs from version 3.1 of the MiBIG database which were not included in version 2 (12) with randomly selected proteins from the representative genomes of GTDB (23), thus ensuring that we did not introduce any previously unidentified BGCs into the benchmark; and (iii) a synthetic metagenome, derived from fragmenting the synthetic isolate genome used in (ii), into contig lengths following the distribution of metagenomic assemblies from a range of environments (see Methods and Supplementary Figure 3).

SanntiS achieved the highest AUCPR among all the methods evaluated for all three tested datasets. Overall, the results of the benchmark show that SanntiS outperforms other methods in genomic and metagenomic data. This high recall rate, alongside high precision, is also reflected in the F1 score achieved by SanntiS. On the “9 genomes dataset” and utilising cluster-based metrics, AUCPR was significantly higher for SanntiS (Figure 1a), followed by antiSMASH, GECCO, and deepBGC. The average-based metrics showed a decrease in precision for all methods (Supplementary figure 4a). While this was particularly pronounced in antiSMASH, it was a common challenge for all methods, reflecting the inherent difficulties in delineating the precise boundaries of the BGCs.

The synthetic isolate provides a greater certainty in the ground truth of BGC delineation. In this benchmark, SanntiS, antiSMASH, and GECCO performed similarly when using cluster-based metrics, with SanntiS displaying marginally better performance (Figure 1b). All three tools showed very high precision, but importantly SanntiS had superior qualitative BGC detection with average-based metrics (Supplementary figure 4b). Performance across different BGC classes further revealed SanntiS’s superiority in macro-average recall (Figure 1c), with a value of 0.763, compared to 0.753, 0.723, 0.624 of deepBGC, GECCO and antiSMASH, respectively. While there may be differences among the benchmarked tools in terms of eukaryotic BGC composition in their training datasets or their prediction scope, the performance in BGC predictions remains consistent, even for classes with a high presence of eukaryotic BGCs, as evidenced in the synthetic isolate dataset (see Figure 5 footnote). Collectively, these results indicate that SanntiS is an effective tool for broad BGC detection.

When analysing the synthetic metagenome, precision remained consistent for all tools, but overall recall diminished relative to the synthetic isolate. SanntiS’s recall was the highest (0.449), but the common reduction in recall highlights the challenge of BGC detection in fragmented metagenomic datasets. The F1 score was also highest for SanntiS (0.615), followed by antiSMASH, GECCO, and deepBGC (0.608, 0.435, and 0.233, respectively).

In summary, our findings, based on diverse benchmarking results, demonstrate that SanntiS consistently performs well on BGC detection across diverse datasets and metrics. Its flexibility, adaptability, and balanced performance emphasise its promise as a method for advancing the exploration of microbial communities.

### SanntiS detects BGCs undetected by other methods

Encouraged by the benchmarking results, we applied SanntiS to isolate genomes to understand the diversity of the predictions compared to the same BGC detection methods evaluated in the benchmarking. To do so we predicted BGCs in the species representative genomes of GTDB (release 86). The results of our analysis indicate that SanntiS produces results highly consistent with other methods as indicated by the upset plot (Figure 2a), which shows a high level of overlap between SanntiS results and those produced by other methods. However, SanntiS uniquely identified 46, 813 BGC predictions.

**Figure 2:**
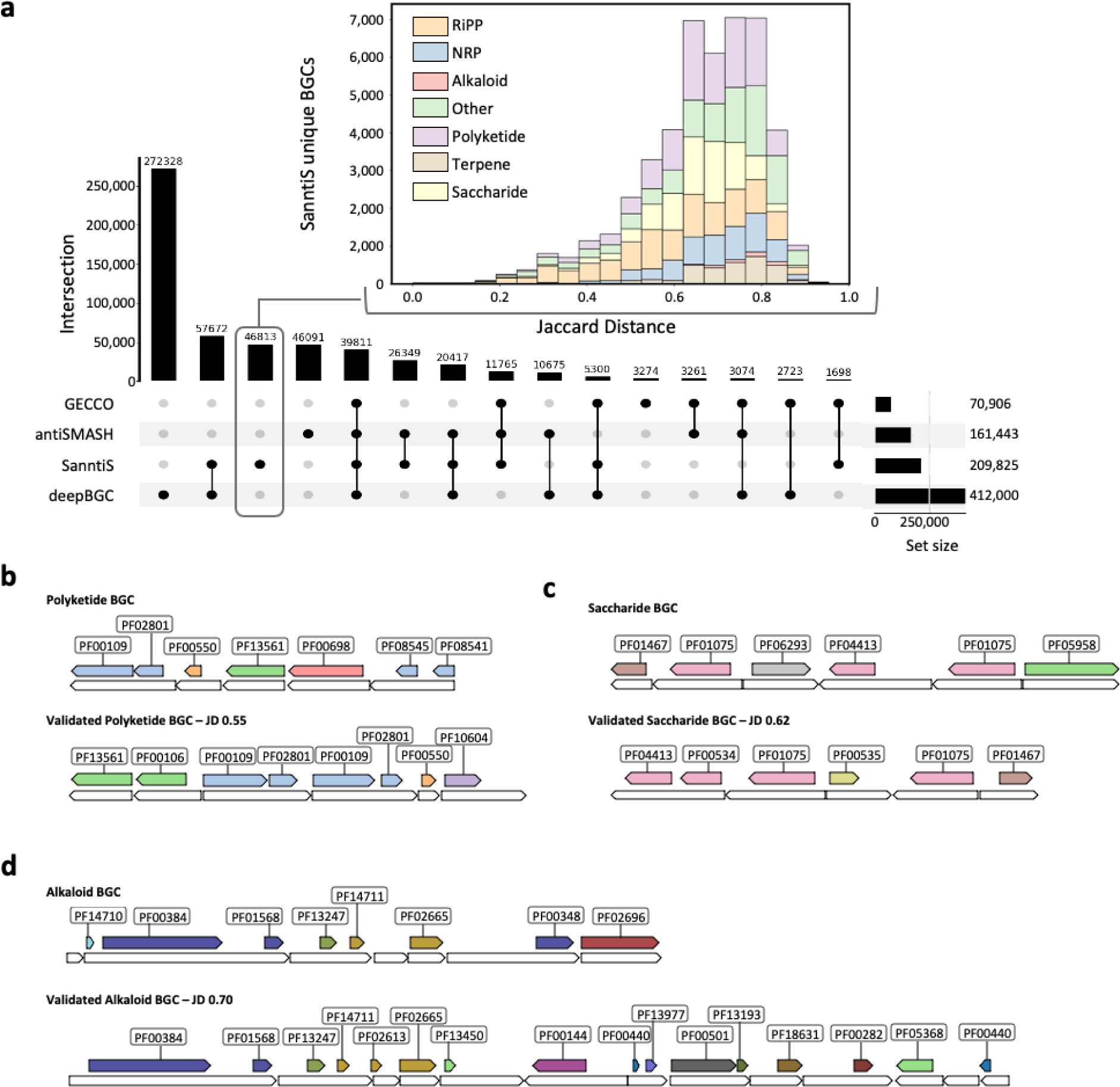
Putative BGCs detected only by SanntiS. (a) Upset plot showing the intersection of BGC regions predicted by each method. Histogram of Jaccard Distance from BGC regions predicted only by SanntiS to the nearest BGC in MiBIG DB. (b, c, and d) Gene architecture of BGCs from the GTDB representatives found only by SanntiS (top) and their nearest MiBIG BGC by Jaccard distance (bottom). White/empty elements indicate proteins. Coloured elements indicate Pfam domains (coloured by Pfam clan). Labelled boxes indicate the Pfam domain accession. (b) Putative polyketide compared to BGC0000228. (c) Putative saccharide compared to BGC0000779. (d) Putative alkaloid compared to BGC0001137.

To gain further insights into the novel 46, 813 BGC predictions produced by SanntiS (insert panel in Figure 2a), we calculated the Jaccard distance (JD) between the BGCs predicted by SanntiS and the BGCs found in MiBIG (version 3.1). As BGCs can undergo rearrangement, the JD was based on the Pfam domain composition, with smaller JD values indicating a closer similarity between the prediction and MiBIG. This approach indicated that the more dissimilar predictions came predominantly from the BGC class Polyketide, while more similar predictions were typically found for RiPP. The BGCs classified as Other were consistently found across various dissimilarities. Overall, the median JD was 0.66, which was within the expected range, as more similar BGCs (with smaller JD scores) are more likely to have been detected by the other methods, and hence not represent a unique SanntiS prediction.

While the JD scores provide an overall impression of the predictions provided by SanntiS compared to known BGCs, we next investigated how the structure of the BGC compared to the nearest known BGC for different classes of BGCs. There were multiple examples across different JD scores and classes where the predicted BGC and a validated BGC shared many similar features (e.g. Figure 2b, c, d). In our detailed analysis of uniquely detected RiPPs — driven by the comparative ease to experimentally validate them compared to other BGC classes — we discovered a particularly interesting putative BGC within the *Brevibacillus laterosporus* genome. This BGC contained an Open Reading Frame (ORF) annotated as a laterosporulin (LS) peptide (PF17861; Figure 3a), a domain known to be associated with antimicrobial activity.

**Figure 3:**
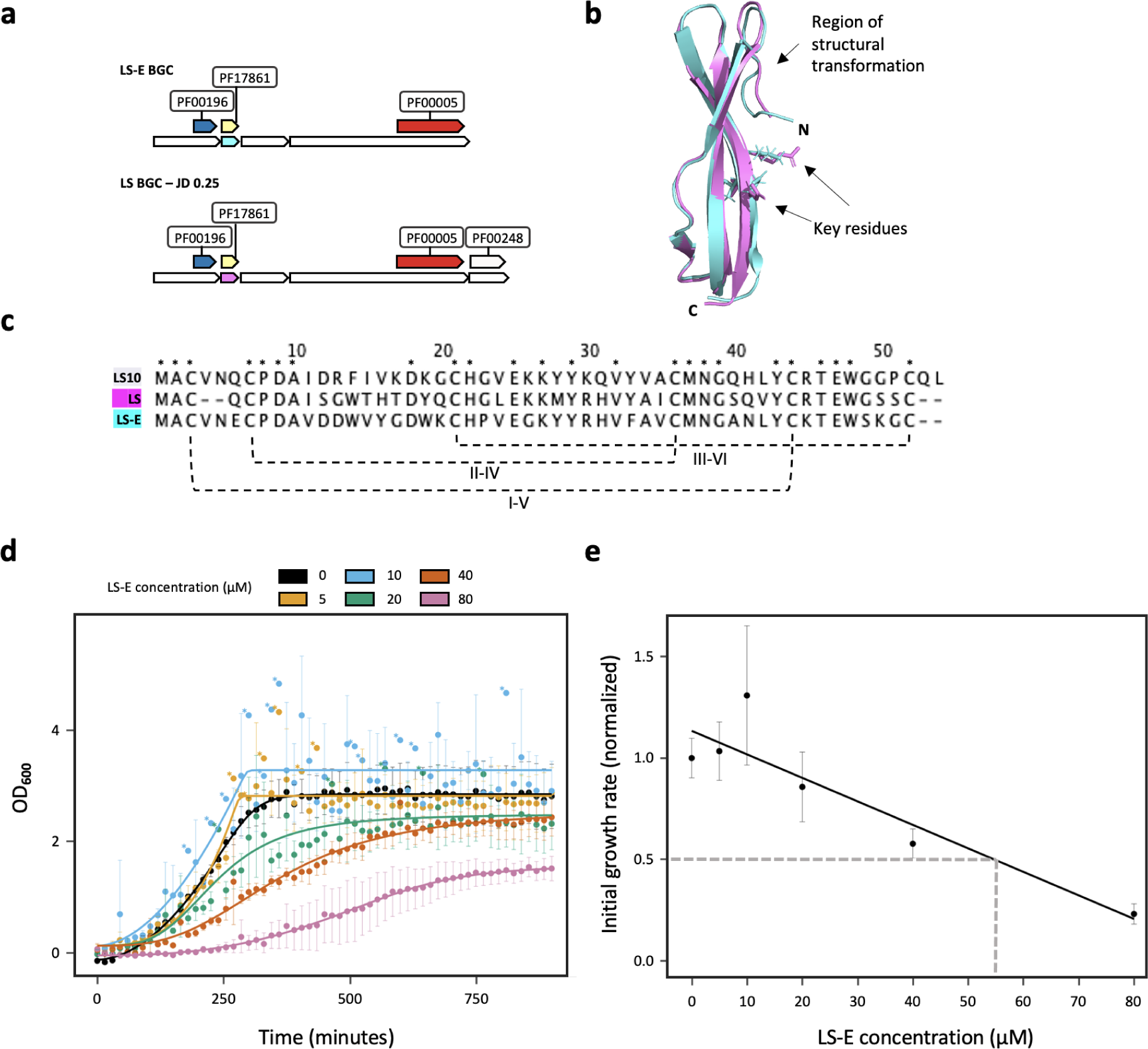
Laterosporulin discovery and experimental validation. (a) Gene architecture of the BGC encoding LS-E compared with the BGC of experimentally validated laterosporulin, LS. Protein domains are coloured according to their Pfam clan. (b) Structure alignment of mature LS (PDB accession 4OZK; magenta) and mature LS-E (predicted structure; cyan). Side chains of key residues in positions 43-45 are shown. (c) Multiple protein sequence alignment of LS-E, LS and LS10. An asterisk (*) indicates positions which have fully conserved residues in this alignment. Disulfide bonds of LS are shown with dashed lines. (d) Growth curves based on OD_600_ measurements of S. aureus cultures incubated with varying concentrations of LS-E. The data are overlaid with curves based on five-parameter Hill equations. Data points are the means of three replicates, with standard deviation indicated by error bars. Data points with very large standard deviations (greater than 1.25 OD_600_ units) are denoted with an asterisk. (e) Initial growth rates vs LS-E concentration. Lag-phase extension, cell killing and growth inhibition are collapsed into a factor representing the average gradient between the zero time point and the sigmoidal curve’s inflection point. All factors are normalised to the negative control (untreated culture). LS-E’s IC_50_ (dashed line) was estimated using linear regression.

As there are no BGCs with this domain in the MiBIG database (and hence our training set), we searched the literature for experimentally validated BGCs with similar characteristics and found two additional examples, which encode for the RiPPs LS (36) and LS10 (37), that had a comparable gene architecture and domain composition. Comparison of the overall BGC organisation revealed an additional ORF in the 3’ end of the BGC that was not covered by the SanntiS prediction. Furthermore, multiple sequence alignment of the predicted laterosporulin (designated LS-E) and the experimentally validated laterosporulins (Figure 3c) indicated that LS-E is more closely related to LS10 than to LS, but the sequences shared many invariant residues in both the leader sequence and the mature peptide (Figure 3c). This high degree of similarity was also found when comparing the determined structure of LS10 (PDB accession 4OZK) and the structural model of the mature LS-E (obtained using AlphaFold 2.0; (38), with a root-mean-square deviation of <0.6Å). The N terminus of mature LS (positions 2 to 7 in the alignment in Figure 3c) has been proposed as a key feature of the sequence for forming a possible trimer (39). This region undergoes a structural transformation from coil (when monomer) to a β-sheet (when trimer), and is stabilised with the β-strand 1 (Figure 3b). Interestingly, LS-E has two insertions (corresponding to positions 4 and 5 in the alignment shown in Figure 3c) in this region that are also present in LS10. LS-E also contains a Glutamine to Glutamic acid substitution at position 6. Another key feature associated with trimer formation is salt bridge interactions between Arginine 45 and Glutamic acid 47. In LS-E Arginine 45 is substituted with Lysine, a conservative substitution. The substitutions at these two positions do not appear to invoke significant structural changes in the position of the side chains based on our structural models (Figure 3b, structure alignment, side chains).

We carried out antimicrobial susceptibility testing of the predicted mature sequence of LS-E. To this end, we incubated the chemically synthesised peptide, at a range of concentrations, with *Staphylococcus aureus* ATCC 25923 (strain MTCC 1430 is a known target of LS and LS10), and assessed the impact on bacterial cell growth via optical density (absorbance) measurements (Figures 3d and 3e). We found that the peptide clearly had antimicrobial activity (IC_50_ = 55 ± 13 μM), even though this activity was somewhat lower than that demonstrated for LS or LS10 against the same bacterial species. This could simply be differences in the experimental assay between laboratories, but could also be that LS-E has a different target specificity or reduced potency. While we have not expressed the complete BGC which may mean that the peptide is not in a native conformation, the antimicrobial activity tests on the chemically synthesised peptide corresponding to LS-E provided additional functional evidence of the novel BGC. Our detailed analysis of the unique RiPP identified by SanntiS against the experimentally validated BGCs with analogous gene architecture, domain makeup, sequence and structural similarity offers compelling evidence supporting its validity. Further experiments involving the expression and testing of the entire BGC would provide more definitive insights into its natural antimicrobial capabilities.

### A collection of BGC predictions from metagenomic samples

To date, the majority of experimentally validated BGCs have been confined to a limited number of taxonomic clades (32 classes in MiBIG 3.1), which may not represent the true biodiversity of BGCs. This hypothesis is substantiated by the BGC predictions on representative genomes in GTDB, which are indicative of more diverse BGC distribution. In light of the positive performance of SanntiS on simulated metagenomic datasets and the broader sampling of microbes enabled by culture-independent approaches, we further explored the BGC repertoire encoded in metagenomes. Application of SanntiS to 33, 924 public metagenomic assemblies from MGnify, encompassing a wide array of ecologically and geographically distinct samples, unveiled 1, 901, 123 putative BGC regions, spanning all known classes (Figure 4a, class panel).

**Figure 4:**
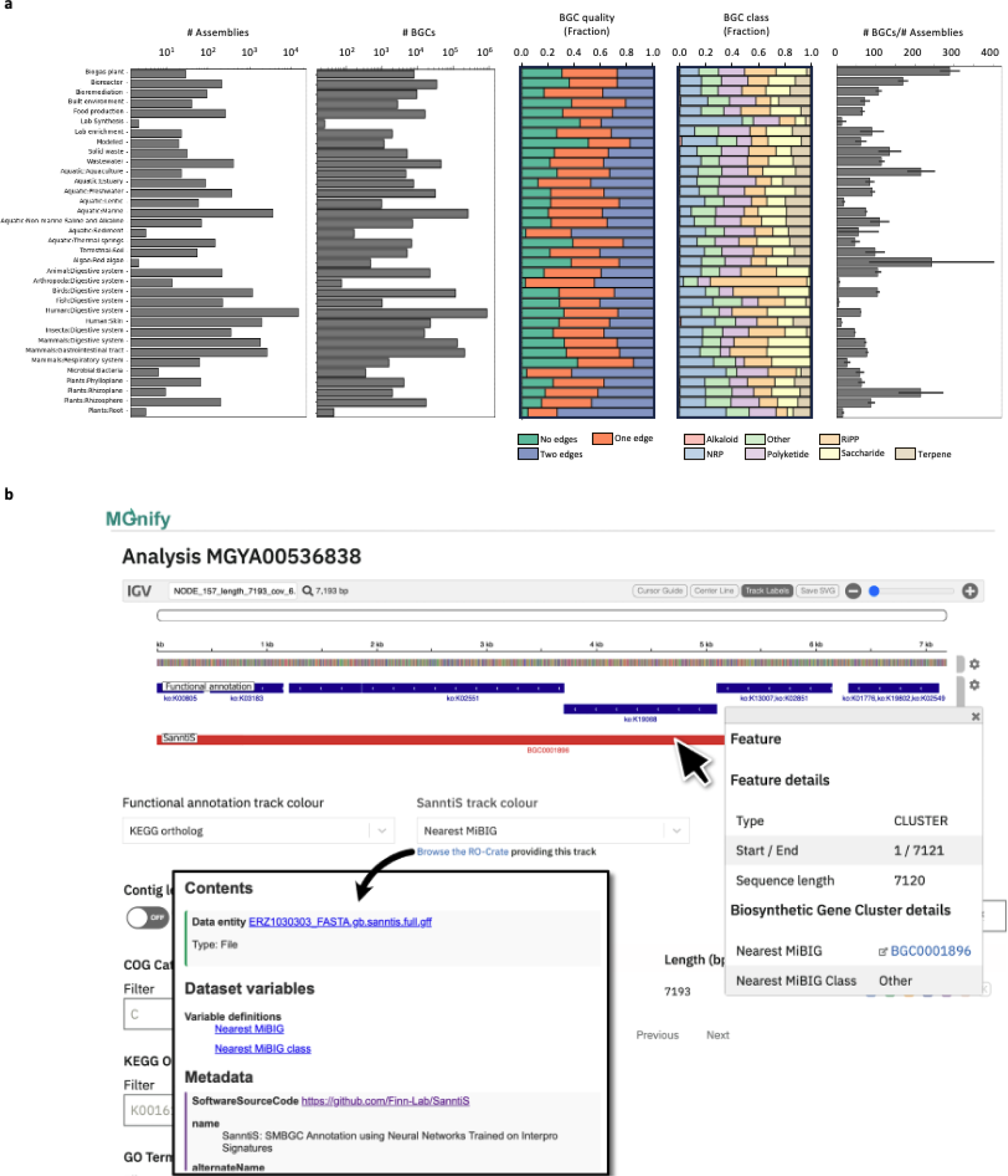
BGC predictions in MGnify assemblies. (a) distinct aspects of BGCs in relation to different biomes. Panels from left to right: Bar plot illustrating the number of assemblies across different biomes; Bar plot presenting the number of BGCs identified in each biome; Bar plot displaying the fraction of BGCs by quality within each biome. Quality is categorised as No edge if BGCs fully contained within a contig, One edge are those overlapping one boundary, and Two edges are those spanning both boundaries; Bar plot depicting the distribution of BGC classes within each biome; Panel 5: Bar plot highlighting the average number of BGCs per assembly across biomes with standard error bars indicating variability. (b) Screenshot of MGnify Contig Browser showing in red a region detected as a BGC by SanntiS. Data associated with the prediction is shown in the feature box. The screenshot pertains to assembly accession ERZ1030303 and contig accession NODE_157_length_7193_cov_6.278509.

Exploring the BGC distribution across various biomes, highlighted the human digestive system, marine environments, mammal digestive system, and birds digestive system as the environments with the highest numbers of predicted BGCs. This observation aligns with the abundant assemblies represented by these biomes. Intriguingly, the human skin biome, despite ranking fourth in terms of assembly count, registered a notably low average number of BGCs per assembly at 11.8, mirroring its comparatively reduced microbial diversity (40). Engineered environments like biogas plants, bioreactors, and solid waste, characterised by high biomass concentrations, emerged as BGC hotspots, with an average of 289.5, 169.3, and 135.1 BGCs per assembly, respectively. This phenomenon corroborates prior studies linking such environments to rampant horizontal gene transfers, encompassing BGCs for polyketides and NRPs (41, 42). Rhizoplane and red algae biomes also presented elevated average BGCs per assembly. However, the limited number of assemblies and significant standard error mean that it would be inappropriate to draw concrete conclusions concerning these biomes.

BGC class distribution across biomes was largely consistent, except in the ‘Lab synthesis’ and ‘Arthropoda digestive system’ biomes. The former exhibited a low fraction of complete BGCs, while the latter displayed a limited number of assemblies. Interestingly, a high proportion of BGCs at the contig edges were RiPPs (shorter BGCs), while a lower proportion favoured extended classes like NRPs and Polyketides. These disparities could, conceivably, be attributed more to assembly limitations than to inherent biome characteristics.

An exemplary advantage of our BGC prediction dataset lies in its integration potential with other analyses furnished by MGnify (Figure 4b), especially the functional annotations and taxonomic profiles, as well as the metadata associated with these datasets. This can render a multidimensional and enriched perspective, facilitating deeper insights into the datasets.

Conclusively, these results highlight a staggering reservoir of BGC predictions across diverse environments, offering valuable insights into the ecological context of BGCs, as well as an otherwise untapped source of new natural products.

### Uncovering the diversity and novelty of BGCs in metagenomic assemblies

Metagenomic datasets are starting to provide access to the vast uncultured microbial diversity, and leveraging this data resource can revolutionise our understanding of BGC diversity. To demonstrate the novelty of BGC predictions found in metagenomics datasets, we compared our BGC predictions from MGnify (denoted as MGnify-SanntiS) to predictions performed on genomic assemblies of bacteria and archaea present in RefSeq (29) using SanntiS. This analysis discerned 2, 711, 361 BGCs across 293, 506 isolate genomes, subsequently termed the RefSeq-SanntiS dataset. For the sake of completeness, we also compared the 2, 502 experimentally validated BGCs from MiBIG DB version 3.1.

Although the RefSeq-SanntiS dataset contained a higher total number of detected BGCs (1.9M for MGnify-SanntiS vs. 2.7M for RefSeq-SanntiS, as visualised in Figure 5a), the BGC class distribution was comparable. Intriguingly, we observed a notable increase in the proportion of NRPs in the RefSeq-SanntiS dataset compared to the MGnify-SanntiS dataset (27% and 15%, respectively). Conversely, RiPPs were proportionally higher in the MGnify-SanntiS dataset, compared to the RefSeq-SanntiS (23% and 12%, respectively). Such disparities are likely to stem from the challenges associated with assembling sufficiently long sequences from metagenomic data that harbour repetitive regions, as found in NRPs.

**Figure 5:**
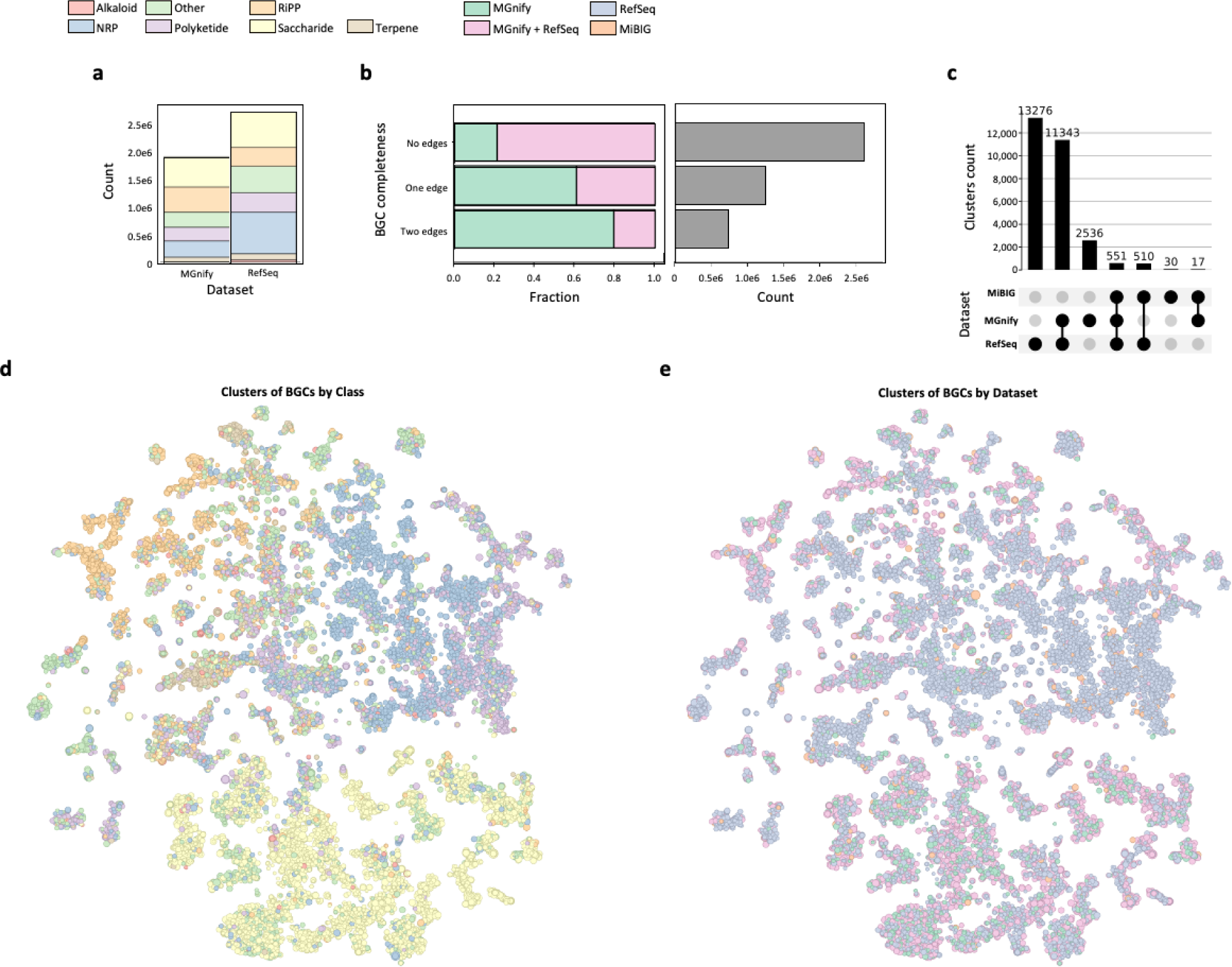
Comparative analyses of BGCs between metagenomic (MGnify-SanntiS) and genomic (RefSeq-SanntiS) datasets. (a) Stacked bar plot depicting the total count of BGCs in each dataset (MGnify-SanntiS vs. RefSeq-SanntiS). Each colour segment represents a distinct BGC class. (b)Categorization of BGCs based on their relative position to contig boundaries. The first panel presents the proportion of BGCs from each dataset across the three defined categories. The second panel exhibits the absolute count of BGCs for each category. (c) Upset plot illustrating the intersections between datasets (MGnify-SanntiS, RefSeq-SanntiS, and MiBIG). The plot showcases shared and unique clusters among the three datasets. (d) t-SNE plot of BGC clusters, coloured according to BGC class, highlighting the distribution and diversity of the different classes. (e) t-SNE plot of BGC clusters, coloured based on dataset origin, emphasising the intersection of clusters across the datasets.

We categorised BGC predictions based on their position relative to contig boundaries: No edges, where the BGC prediction was fully contained within a contig; One edge, if the prediction ended at one contig boundary; Two edges, where the prediction spanned the whole contig (with the presumption that these BGCs may be incomplete and therefore lower quality). Unsurprisingly, the majority of No edges BGCs originated from RefSeq, and MGnify-SanntiS predictions increased with the number of contig edges, reflecting the fragmented nature of metagenomic assemblies (Figure 5b).

To demonstrate the difference in BGC predictions between the MGnify-SanntiS and RefSeq-SanntiS datasets, we clustered the complete BGC predictions (i.e. no contig edges) based on domain composition to prevent bias that may be introduced from incomplete predictions. Of the 28, 263 resulting clusters, 9% (2, 536) were exclusive to the MGnify-SanntiS dataset. While 11, 343 clusters were shared between MGnify-SanntiS and RefSeq-SanntiS datasets, 13, 276 clusters were confined to the RefSeq-SanntiS (Figure 5c). The experimentally validated BGCs from MiBIG were found in 1, 108 clusters, with 568 of these also containing members from MGnify-SanntiS. Overall, this reflects an increase in the number of clusters, highlighting the novelty of BGCs found in metagenomics datasets, such as those contained in MGnfy, as well as isolate genomes having a different distribution to environmental samples.

To understand the distribution of different BGC classes coming from the two datasets, we coloured the clusters by BGC class (Figure 5d) and data source (Figure 5e). Comparing the two figures shows that, despite the relatively even counts of most BGC classes (Figure 5a), the domain diversity of some BGC classes (and hence number of clusters) was substantially higher than others, especially Saccharides. This observed pattern agrees with a similar analysis performed on isolate genomes by Cimermancic *et al*. (16), which also revealed a pronounced prevalence of Saccharides. Interestingly, clusters exclusive to MGnify-SanntiS were dispersed across all classes.

## Conclusion

In this study, we introduced SanntiS, a novel machine learning-based approach for identifying biosynthetic gene clusters (BGCs) in genomic and metagenomic sequences. Our results indicate that SanntiS surpasses existing methods in terms of performance, recall, and applicability. We believe that the superior efficacy stems from the application of range events-specific training. These techniques effectively capture the sequential nature of BGCs and address the challenges of length imbalance across classes. While such techniques have been applied in domains like sound event-detection and time-series applications, to our knowledge this is the first time they have been used for biological sequence annotation assessments.

While we have only been able to perform experimental validation on a single bacteriocin, this was selected as it was predicted only by SanntiS and was absent from the underlying training set. Although the potency of the LS-E peptide is relatively weak against the tested strain, further testing of LS-E against different bacterial strains may identify more sensitive strains to this RiPP, while expression *in vivo* may produce a topology with enhanced antimicrobial activities. Nevertheless, this underscores the potential for SanntiS to expand the range of known natural products.

The large-scale application of SanntiS to the metagenomic assemblies produced 1.9 million BGC predictions. This collection represents one of the most expansive compilations of BGCs derived from metagenomic datasets to date. Other significant endeavours (8, 43) have predominantly focused on predictions from Metagenome Assembled Genomes (MAGs), but these neglect the plethora of contigs from lower abundant organisms that fail the MAG generation process, as well as plasmids. Furthermore, this substantial resource holds much promise for further characterisation via informatics and experimental validation. As we continue to explore the vast metabolic potential of microbial communities, tools and datasets like these will be fundamental in bridging knowledge gaps and catalysing innovations in natural product discovery and ecological research.

## Methods

### Open reading frame (ORF) identification and functional annotation

Prodigal v2.6.3 (21) was used for ORF identification and protein translation for all the DNA sequences utilised in this work. The resulting proteins were then functionally annotated using a subset of the member databases in InterProScan v5.52-86.0 (Pfam, TIGRFAM, PRINTS, ProSitePatterns, Gene3D) (22) and a set of profile hidden Markov models (HMMs) from Pfam version 34.0, which had not yet been included in InterProScan. This subset of member databases was selected based on those employed in MGnify to strike a balance between the speed of annotations and coverage.

### Training and validation datasets

A training and validation dataset of known BGCs was constructed using the nucleotide sequences from the MiBIG database version 2.0 (20), taking BGCs from all classes. To remove redundancy within this dataset, and given the known rearrangements of genes within a BGC, the MiBIG BGC sequences were clustered based on the Jaccard similarity score of InterProScan entries (cut-off threshold of 0.95), with the longest sequence retained as a representative. The same approach was employed to remove any MiBIG sequences found in the training set that were similar to the test datasets (described below). In total, 1, 858 BGC (positive) entities were used in the training and validation datasets.

For the development of a negative dataset, representative genomes from GTDB (release 86; (23) were used as a source of non-BGC regions. Potential BGC sequences were identified and removed using antiSMASH v5.1. To avoid imbalance between the positive and negative datasets, the non-BGC regions were subsampled by randomly selecting 5, 574 chunks of consecutive ORFs (with length sampled from a normal random distribution with the same mean and standard deviation as the positive dataset).

A synthetic contig was also developed to train the models. The contig was created by first duplicating the positive entities to increase the robustness of the training. The BGC and non-BGC entities were then shuffled to produce a synthetic contig for model training. To assess the performance of the models, a cross-validation approach was used that involved five independent and stratified splits of positive classes for validation, following a similar methodology to the training contig, but excluding the duplication of positive entities. The synthetic contigs generated from these procedures were utilised to train and validate the performance of the models.

### Test datasets

To evaluate the method and compare performance with other BGC finding methods, 3 different datasets were used: (i) the “9 genomes ”, originally used in the work of (16), and revisited and recollected by (18), with manually annotated BGC and non-BGC regions; (ii) a test dataset of newly published BGC sequences not found in MiBIG version 2, and independent of the training and validation datasets.

The construction of this dataset was initiated by determining positive regions from the difference between MiBIG versions 3.1 and 2.0 (12) . To avoid increasing complexity, the class of each of these BGCs was assigned by keeping only the first class (when more than one was in the annotation) in alphabetical order. Using the GTDB release v214.1, dated June 9th 2023, we extracted lineage information for the representative genomes. From the available 202 phyla, 70 were randomly selected. For each of these phyla, a representative genome was sampled. The selected genomes were processed with antiSMASH v6.1, and coding sequences (CDS) that were identified as core BGC were excluded to produce non-BGC regions. These regions were then shuffled and divided into 70 different contigs for computational efficiency. Concurrently, positive regions were shuffled and split into 70 distinct lists (to align with the earlier step), and were inserted at random points within the corresponding non-BGC regions, creating what we term the “synthetic isolate genome ”. The results of this assembly are available in the table located at Supplementary table 1 and in Supplementary FASTA 1.

To examine the performance of the BGC detection methods when executed on metagenomic assemblies, a “synthetic metagenomic assembly ” was created. For this assembly, a random selection of 1, 000 MGnify assemblies, originating from a variety of environments, was made. The primary objective was to analyse the length distribution of the contigs they housed. With 5, 135, 756 contigs to observe, we calculated the probability density function (pdf) using the gaussian_kde function provided by Python Scipy. This model then served as the basis for sampling the distribution. The synthetic contigs were fragmented according to this distribution. The entire procedure was replicated three times to generate three distinct sets of data. The specific details of the fragmentation are found in the headers of the FASTA files of Supplementary FASTA 2.

### Feature matrices construction

Feature matrices appropriate for model training and prediction were constructed from a linear sequence of annotation entries, ordered by their start position. Subsequently, the domains were one-hot encoded.

### Model training and BGC prediction implementation

An artificial neural network (ANN) model consisting of 4 layers was built using TensorFlow v2.4.1 (24). First, an embedding layer was employed to represent each functional annotation as a vector of 124 features. Second, a one-dimensional Convolutional (Conv1D) layer of 64 filters and a kernel size of 4 was used. Third, a bidirectional long-short term memory (BiLSTM) layer of 32 units was implemented, which conserved the original input dimensions, and a dropout layer of 0.5 was connected to a single-unit dense layer, activated with a sigmoid function. The training used the minibatch setup and applied the Adam optimizer with the learning rate dropping by a factor of 0.1 every 5 epochs of not improving the loss function on the validation dataset. A custom implementation of RLF with a gamma factor of 2.0 was used as the loss function. The training was performed using sequences of 200 timesteps.

The BGC detection process involved predicting the probability of each of the linearized annotation entities and normalising it by retaining the entity with the maximum value for each ORF. Then, given a score threshold, the regions of the sequences above that threshold were identified as BGCs. A Multiclass OneVsRest Random Forest classifier was also trained (using default parameters of scikit-learn v0.24 (25) for the 7 MiBIG classes plus a non-BGC class, using the same feature matrices that were used for the ANN model. This classifier served as a post-processing filter, where predicted regions not classified as one of the 7 MiBIG classes were dropped.

### Validation of SanntiS performance and comparison with other methods

Performance evaluation metrics, such as recall, precision, F1, and F2 score, were utilised. To avoid known problems of evaluations using short segments as units (26), two different approaches were employed for deriving score metrics.

Cluster-Based. In this approach, the entire BGC region was used as the unit of measure, demonstrating the method’s ability to detect BGCs but omitting the quality of the predictions. A true positive (TP) was a predicted BGC region overlapping a BGC region in the reference, a false positive (FP) was a predicted BGC region not overlapping any BGC in the reference, and a false negative (FN) was a BGC region in the reference not overlapping any predicted BGC.

Average metrics. Micro-averaging of intermediate statistics was implemented in this approach, which is better suited for range-based anomalies (like BGCs) as it can demonstrate the quality of the predictions and avoid biases induced by length differences, giving equal importance to all BGCs (irrespective of their length). For each BGC region in the reference, RecallT was calculated as the number of ORFs predicted in a BGC region divided by the ORFs in the reference region, with the average of this metric then being taken. For each predicted BGC, PrecisionT was calculated as the number of ORFs that were in a BGC in the reference divided by the total number of predicted ORFs, followed by obtaining the average of the individual precisions (27).

Validation of the method and comparison of SanntiS v0.9.3 with antiSMASH v6.1.1, deepBGC v0.1.30, and GECCO v0.9.8 in real genomic contexts was performed by running the methods at different score/thresholds (from 0 to 1 with a step of 0.005, for deepBGC, GECCO, and SanntiS. Strict, relaxed, and loose modes were used for antiSMASH) on every dataset, with all other parameters set to default. Recall metrics for individual BGC classes were also calculated. BGCs in the “9 genomes ” dataset were classified by obtaining the Jaccard distance (JD) of the BGCs to those in MiBIG, with JD as given by

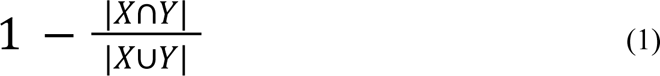

where X are the Pfam domains in one region and Y are the Pfam domains in the other.

To evaluate the performance of each method in an independent validation dataset and to assess the differences in detection between a long contig genome assembly and a metagenomic assembly, evaluation metrics for the synthetic isolate genome dataset were derived using the default parameters of each method and the classes assigned by the MiBIG curators. For the prediction in the synthetic metagenomic assembly, the Prodigal “meta mode ” was used in antiSMASH (with a minimal length of 500 bp) and deepBGC. For the synthetic metagenomic assembly, a true positive (TP) in the Cluster-based metrics was considered the detection of at least one CDS of a true BGC, irrespective of the number of contigs that particular BGC spans.

### BGC prediction in GTDB representative genomes

The 31, 910 GTDB representative genomes were downloaded, and BGC regions were predicted using SanntiS, antiSMASH, deepBGC, and GECCO with default parameters. Overlapping predictions were then identified, and regions called only by SanntiS were selected. For every region, the Jaccard distance (JD) to the BGCs in MiBIG was calculated, and the assigned BGC class corresponded to that of the nearest BGC in MiBIG. Some randomly selected BGC regions called only by SanntiS were manually examined and compared with those in MiBIG. For one of these cases, which showed strong evidence of being a true BGC, a multiple sequence alignment of the putative structural protein with those of similar known BGCs was constructed. AlphaFold version 2.0, was used to predict the structure of the putative structural protein.

### Experimental validation of predicted BGC

Chemically synthesised protein LS-E was ordered from WuXi AppTec (TFA salt, 84 % purity), resuspended in Milli-Q water and filter-sterilised (PVDF; Millex SLGV004SL). Antimicrobial peptide Assay Buffer (AAB; 0.01 % (v/v) acetic acid, 0.2 % (w/v) BSA) was prepared using Milli-Q water, glacial acetic acid (Merck 27225) and BSA powder (Sigma A9418), and filter-sterilised (PES; Millipore SLGP033RS).

Streaked LB-Lennox agar plates (1.6 %; Formedium AGR10, Formedium LBX0102) of *S. aureus* (ATCC 25923) were incubated at 37 C overnight, then individual colonies were picked for overnight culture in liquid LB-Miller medium (Formedium LMM0102). This overnight culture was diluted to optical density (OD_600_) 0.125 in Mueller-Hinton broth (Millipore 70192) before adding an equal volume of LS stock solution, then establishing a two-fold titration series by repeated dilution in OD 0.125 bacterial culture in a microtitre plate (Eppendorf 0030602102), in triplicate. Plates were sealed (SLS PCR0548) and per-well absorbance at 600 nm was measured *via* spectrophotometer (FLUOstar Omega) to monitor bacterial growth during an overnight incubation at 37 C. All plasticware used was polypropylene.

To fit the data, the following starting parameters were used: effect coefficient, E = 250; Hill coefficient, H = 10; bottom asymptote, B = 0; top asymptote, T = 3; symmetry coefficient, S = 1. A linear approximation to the Hill equation (with fixed B, T and S) was first used to obtain estimates for E and H, which then replaced their corresponding starting parameters for a non-linear model (five-parameter Hill equation; y = B + (T – B)/(1 + (2^(1/S)^ - 1)((E/x)^H^))^S^), constrained as follows: 150<E<750; 3<H<100; -0.125<B<0.125; 1.25<T<3.5; 0<S<1. The average initial growth rate (dy/dx) for each concentration was determined by dividing the half-maximal OD_600_ by E, then normalised to the initial growth rate measured for untreated culture. The IC_50_ for LS-E was estimated by fitting these initial growth rates via linear regression. Data analysis and plot generation was performed in R.

### BGC prediction in genomic and metagenomic assemblies

Contigs longer than 3, 000 base pairs were extracted from 33, 924 publicly accessible metagenomic assemblies, provided by the MGnify resource (28). BGC regions were predicted within the contigs using SanntiS, with default parameters. In a parallel effort, BGCs were predicted on Archaeal and Bacterial genomes present in RefSeq (29) (downloaded on 20th July 2023) using the SanntiS default parameters.

The details regarding these predictions can be found in Supplementary Table 2.

### BGC clustering analysis

Regions from the predictions that did not extend to the edge of a contig were integrated with experimentally validated BGCs from the MiBIG Database. This merged dataset was then used for the subsequent cluster analysis.

Clusters of BGCs based on functional domain composition were identified in the following manner: Regions containing the same set of Pfams were identified and collapsed into one entity (Supplementary Figure 4). Using the Python package NetworkX (https://github.com/networkx/networkx), a similarity network was built where a node represents a set of Pfams, and the weight of the edges is the Dice coefficient (DC), given by

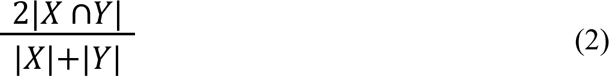

for all pairs in the collection, where |XY| is the longest overlapping string between the two sequences and |X|+|Y| is the sum of both sequences’ lengths. To reduce the number of edges and computational expense, a K Neighbours Graph was built, with a K parameter equal to 14 (30, 31). In total, 570, 511 sets of Pfams were clustered into 33, 905 groups by the Markov Clustering algorithm (32) (The implementation found on https://github.com/GuyAllard/markov_clustering), with an inflation of 1.8. The inflation value was determined as the highest modularity score (33) in an interval from 1.70 to 4.0 with increments of 0.05. Clustering analysis results are in Supplementary Table 3. Note that the bgc_names in this table correspond to those in Supplementary Table 2.

### BGC clusters graph

For each BGC cluster, the similarity to other clusters was calculated as the average DC of all its members against all members of the other cluster. An epsilon-graph with cutoff threshold of 0.45 was built for visualisation purposes. Epsilon value was calculated using the “Elbow-method” heuristic. Embedding of the nodes was performed using the ProNE algorithm with 124 dimensions. Thereafter, a 2 dimensional projection using T-SNE (perplexity=70) with Scikit-learn (25)

## Supporting information

Supplementary FASTA 1

Supplementary FASTA 2

## Data availability

The SanntiS software implementation is accessible as a standalone Python package and can be downloaded via the GitHub repository at https://github.com/Finn-Lab/SanntiS. The supplementary materials contain detailed information on data accessions and resource location, as well as the necessary data for replicating the training dataset to ensure reproducibility. The data employed for testing and validation in this study is available in supplementary data. BGC predictions by SanntiS on metagenomic assemblies are readily downloadable from the MGnify website. Assemblies that were analysed using MGnify’s pipeline version 5 can be visualised using the portal contig viewer.

## Acknowledgements

We extend our deepest gratitude to Olivia Casanueva and Lorna Richardson, both from EMBL-EBI, for their diligent proofreading efforts, which have greatly enhanced the clarity and readability of our work. We particularly appreciate Lorna Richardson for her additional support and insightful contributions throughout the development process.

## Supplementary Information

**Supplementary Table 1**

Table with the specifications of synthetic isolate genome

**Supplementary FASTA 1**

FASTA files of synthetic isolate genomes

**Supplementary FASTA 2**

FASTA files of synthetic metagenomic assembly

**Supplementary Table 2**

Detected BGC regions by SanntiS on RefSeq and MGnify datasets

**Supplementary Table 3**

Clustering analysis results on high quality BGCs

**Supplementary Table 4**

Selected examples of BGCs from the GTDB Representative Genomes detected exclusively by SanntiS.

## Supplementary Figures

**Supplementary Figure 1:**
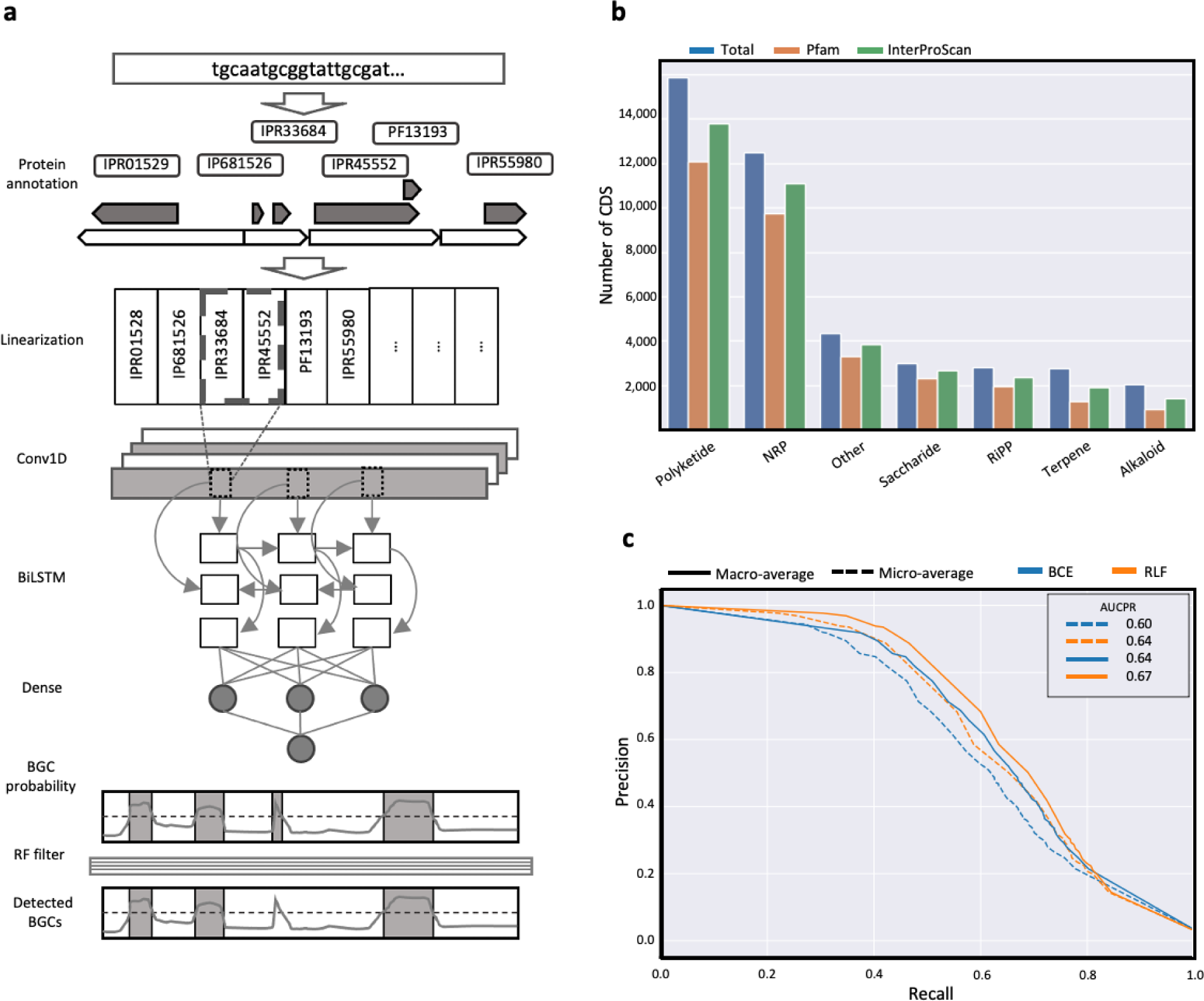
SanntiS data flow and unique features of the method. **(a)** Data flow in SanntiS detection method. Nucleotide sequences are used as input, then proteins are predicted to be functionally annotated. The annotations are linearized to go into an artificial neural network (ANN) with a 1D convolutional (Conv1D) layer and a bidirectional long-short term memory (BiLSTM). Regions with a probability above the threshold are filtered by the RF classifier. **(b)** Proteins per BGC class in MiBIG database and their annotation. Blue, total number of proteins. Orange, proteins annotated by Pfam. Green, proteins annotated by the subset of InterProScan. **(c)** 3×5 fold cross-validation precision-recall curves. Colour indicates the loss function used for training. Standard binary cross-entropy loss function (BCE) and the Robust loss function (RLF). Line style indicates the type of metric used.

**Supplementary Figure 2:**
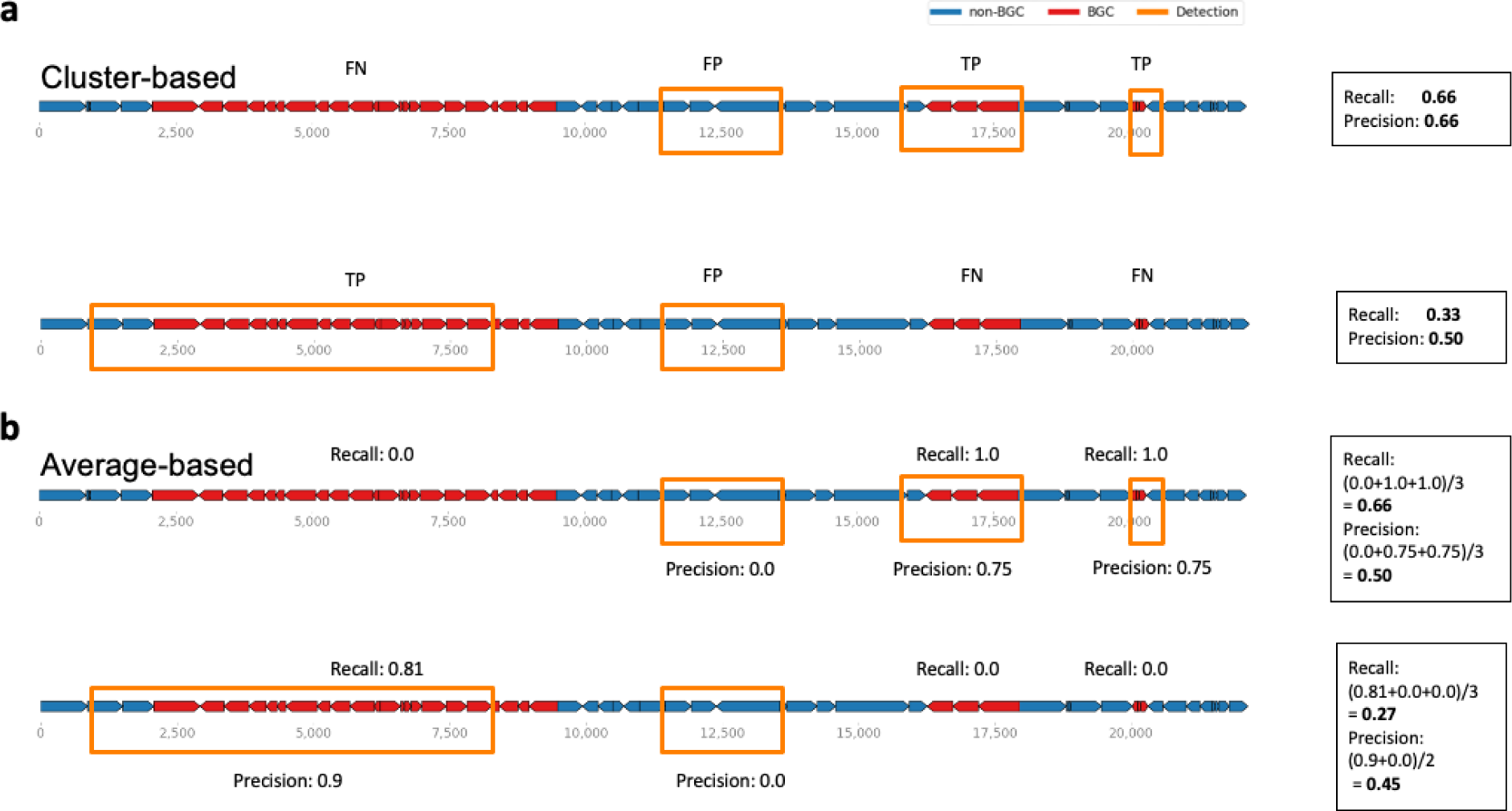
Evaluation metrics. Illustration of the hypothetical DNA fragments with coding sequences (CDS) of ground truth BGCs (red) and non-BGC (blue). Yellow boxes represent the predicted BGC regions. **(a)** demonstrates the calculation of metrics using the cluster-based approach, where the overlap of a predicted region with the ground truth BGC is considered as a true positive (TP) and a predicted region not overlapping any ground truth BGC regions is a false positive (FP). **(b)** average-based approach, where precision and recall are calculated for each predicted BGC and ground truth BGC by dividing the number of TP CDS by the total CDSs in the prediction and BGC, respectively, and then averaging the results across all predicted regions. This approach is utilised to avoid ambiguity caused by unbalanced datasets and uneven length distributions of BGCs.

**Supplementary Figure 3:**
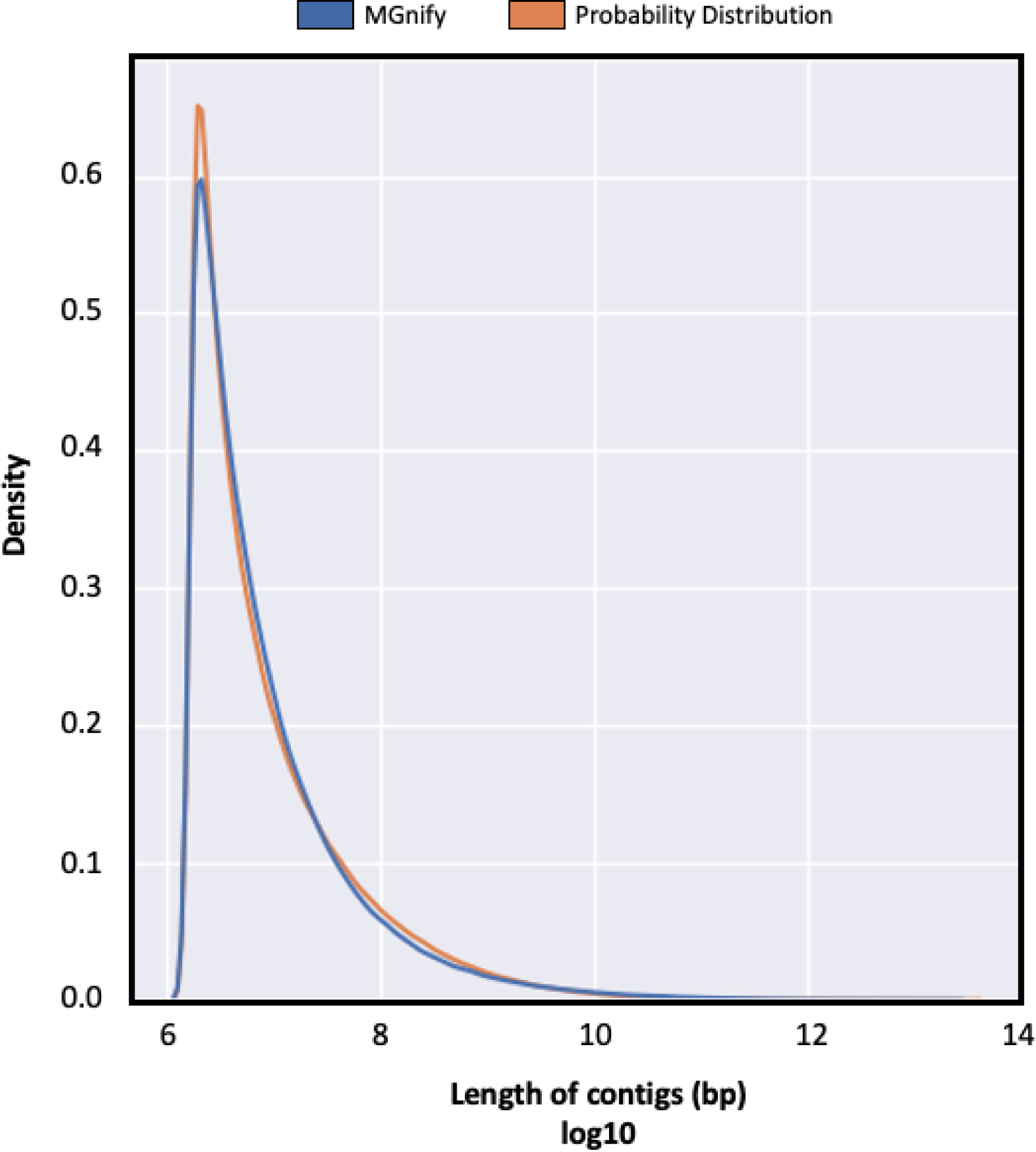
Length distribution of metagenomic contigs, presented as a density plot. The x-axis represents the length of contigs in base pairs on a log10 scale, while the y-axis denotes the density. The blue line represents the observed data from MGnify, which illustrates the distribution of metagenomic assemblies from diverse environments. The orange line shows the modelled random probability distribution, which is used to compare and analyse the relationship between the observed data and the expected distribution. This figure helps to provide insights into the contig length distribution in metagenomic datasets and its potential impact on BGC detection methods.

**Supplementary Figure 4:**
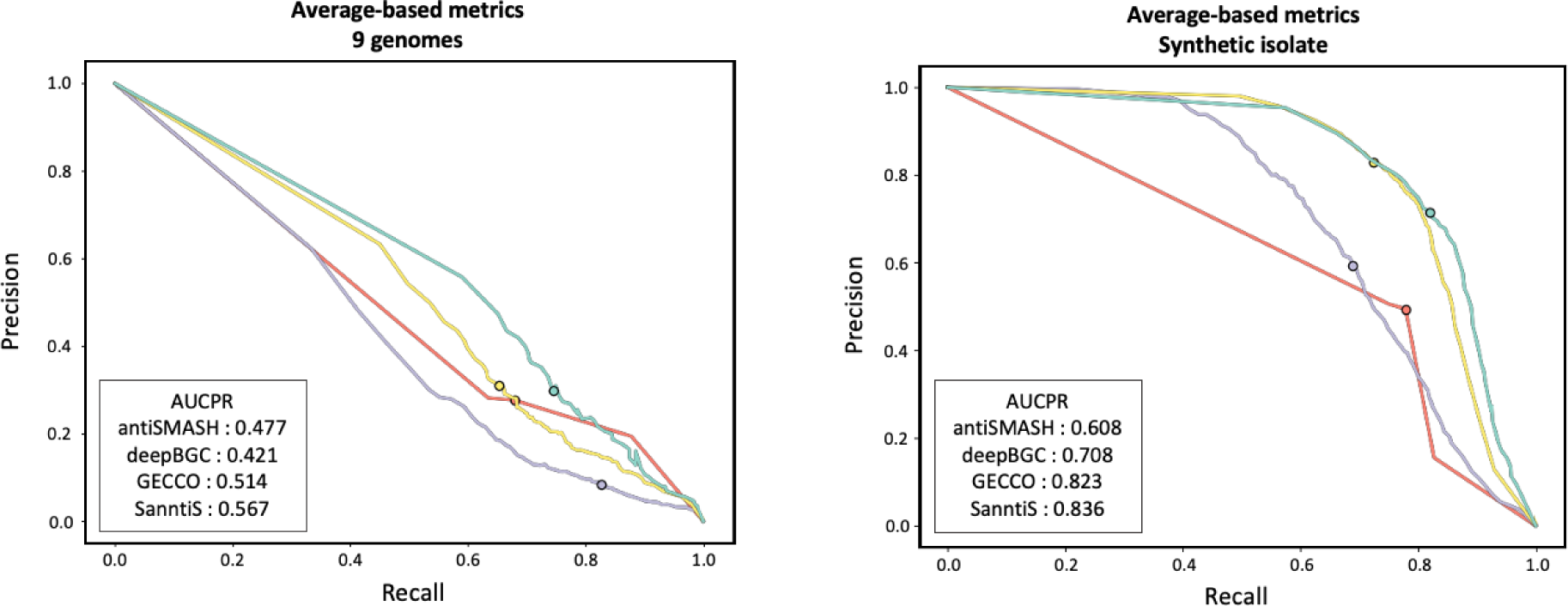
Benchmark results**. (a)** Average-based precision-recall curve of each method at different probability thresholds in 9 genomes. The area under the precision-recall curve (AUCPR) is shown for those methods with the option of changing probability thresholds. **(b)** Average-based precision-recall curve of each method at different probability thresholds in Synthetic isolate. The area under the precision-recall curve (AUCPR) is shown for those methods with the option of changing probability thresholds. **(b)** Average-based precision-recall curve of each method at different probability thresholds in Synthetic isolate. The area under the precision-recall curve (AUCPR) is shown for those methods with the option of changing probability thresholds.

